# A Prism Vote Framework for Individualized Risk Prediction of Traits in Genome-wide Sequencing Data of Multiple Populations

**DOI:** 10.1101/2022.02.02.478767

**Authors:** Xiaoxuan Xia, Rui Sun, Yexian Zhang, Yingying Wei, Qi Li, Marc Ka Chun Chong, William Ka Kei Wu, Benny Chung-Ying Zee, Hua Tang, Maggie Haitian Wang

## Abstract

Multi-population cohorts offer unprecedented opportunities for profiling disease risk in large samples, however, heterogeneous risk effects underlying complex traits across populations make integrative prediction challenging. In this study, we propose a novel Bayesian probability framework, the Prism Vote (PV), to construct risk predictions in heterogeneous genetic data. The PV views the trait of an individual as a composite risk from subpopulations, in which stratum-specific predictors can be formed in data of more homogeneous genetic structure. Since each individual is represented by a composition of subpopulation memberships, the framework enables individualized risk characterization. Simulations demonstrated that the PV framework applied with alternative prediction methods significantly improved prediction accuracy in mixed and admixed populations. The advantage of PV enlarges as the sample size, genetic heterogeneity, and population diversity increase. In two real genome-wide association data consists of multiple populations, we showed that the framework enhanced prediction accuracy of the linear mixed model by up to 12.1% in five-group cross validations. The proposed framework offers a new aspect to analyze individual’s disease risk and improve accuracy for predicting complex traits in genome data.

## Introduction

Genome-wide genetic markers encode a considerable portion of common human traits heritability (1). One attractive application of the susceptible single nucleotide polymorphisms (SNPs) is to construct prediction models for assessing disease risk. Previous association studies have demonstrated that most complex traits possess a polygenic background influenced by collective genetic variants of moderate to small effects (2, 3, 4), as exhibited in the human height (2), bipolar disorder (3), and cancers (4). Furthermore, due to genetic heterogeneity of complex diseases and the intricated genetic architecture, a considerable part of the identified risk predisposition loci does not overlap across populations, making integrative analysis of multiple populations challenging.

Although single population analysis allows accurate risk effect estimation in large samples, such condition is often unavailable for the non-European population. Furthermore, a geographically defined single population may still harbor different degree of diversity in phylogeny (5). Therefore, direct combination of populations may render prediction accuracy because of mixing the genetic architectures (6-9). Several methods were proposed to integrate multi-ethics cohorts. Cai *et al*. improved the polygenic risk score (PRS)-based prediction for a target minority population by estimating the transferrable effect of a common set of SNPs between the target and a larger auxiliary population (10). Coram *et al*. developed a linear mixed model (LMM)-based prediction method for the minority population by incorporating the risk loci from an auxiliary population as a random component (8). These methods improve prediction in the target population through refining the effect size and SNP subsets, therefore, individuals carrying the same allelic variations at these SNPs will be estimated with the same degree of risk.

Alternatively, a dimension of individual identity can be incorporated together with the SNP-dimension to improve risk prediction. The disease risk of a subject can be considered as a composite risk shaded from multiple subpopulation strata, in which stratum-specific genetic risk can be characterized, and the overall risk is integrated according to the subject’s propensity to the strata. Therefore, subjects of identical risk alleles may be predicted with different risks according to one’s subpopulation propensity. Under this framework, named the Prism Vote (PV), the subpopulation can be defined as strata of more homogeneous genetic architecture compared to the joint data; it might be shaped from population stratification or imply subpopulations experiencing similar environmental exposures altering gene-environmental interactions. A Bayesian probability framework is used to integrate the stratum-specific risks and subject’s subpopulation propensity. The effect of PV is demonstrated in three simulation studies and two real genome-wide association datasets on six traits.

## Material and Methods

### The Method Overview

The PV leverages on the genetic heterogeneity and polygenicity nature of complex traits. The detection of trait-influencing markers, thousands of variants with modest effect size, are sensitive to the underlying genetic architecture of data. Stratification of samples may lead to the identification of stratum-specific risk loci and effects. The framework obtains the stratum-wise risk estimates and through modelling the disease of an individual as a composite risk outcome from multi-layer subpopulations, delivers the individualized risk probability of traits.

Suppose the risk of a trait for subject *s* is attributed from multiple risk strata. Let *Y* denote a phenotype of binary outcome, and ***x***_*s*_ is the genotype matrix of subject *s*. The disease probability of the individual can be written as,

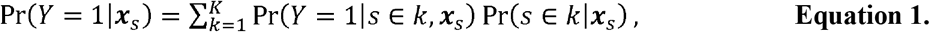

**Equation 1** is referred to as the PV probability of a trait for an individual. It could be generalized to 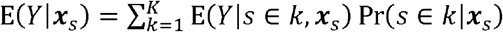 for a continuous *Y*. In the equation, Pr(*Y* = 1|*s* ∈ *k*, ***x***_*s*_) is the disease risk in stratum or subpopulation *k*, to be obtained by a base prediction model; and Pr (*s* ∈ *k*|***x***_*s*_) is the propensity of subject *s* belonging to stratum *k*, calculated by the Bayes theorem:

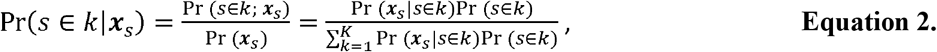

in which Pr(***x***_*s*_|*s* ∈ *k*) is the probability of observing ***x***_*s*_ given subject *s* belongs to stratum *k* ∈ {1, ⋯, *K*}; Pr(*s* ∈ *k*) is estimated by the proportion of the *k*^*th*^ stratum in all samples. **Figure 1** shows a schematic diagram of the PV framework. The term “prism” reflects the interpretation that an individual’s disease risk is decomposed into a spectrum of risk distributions by population strata.

**Figure 1.**
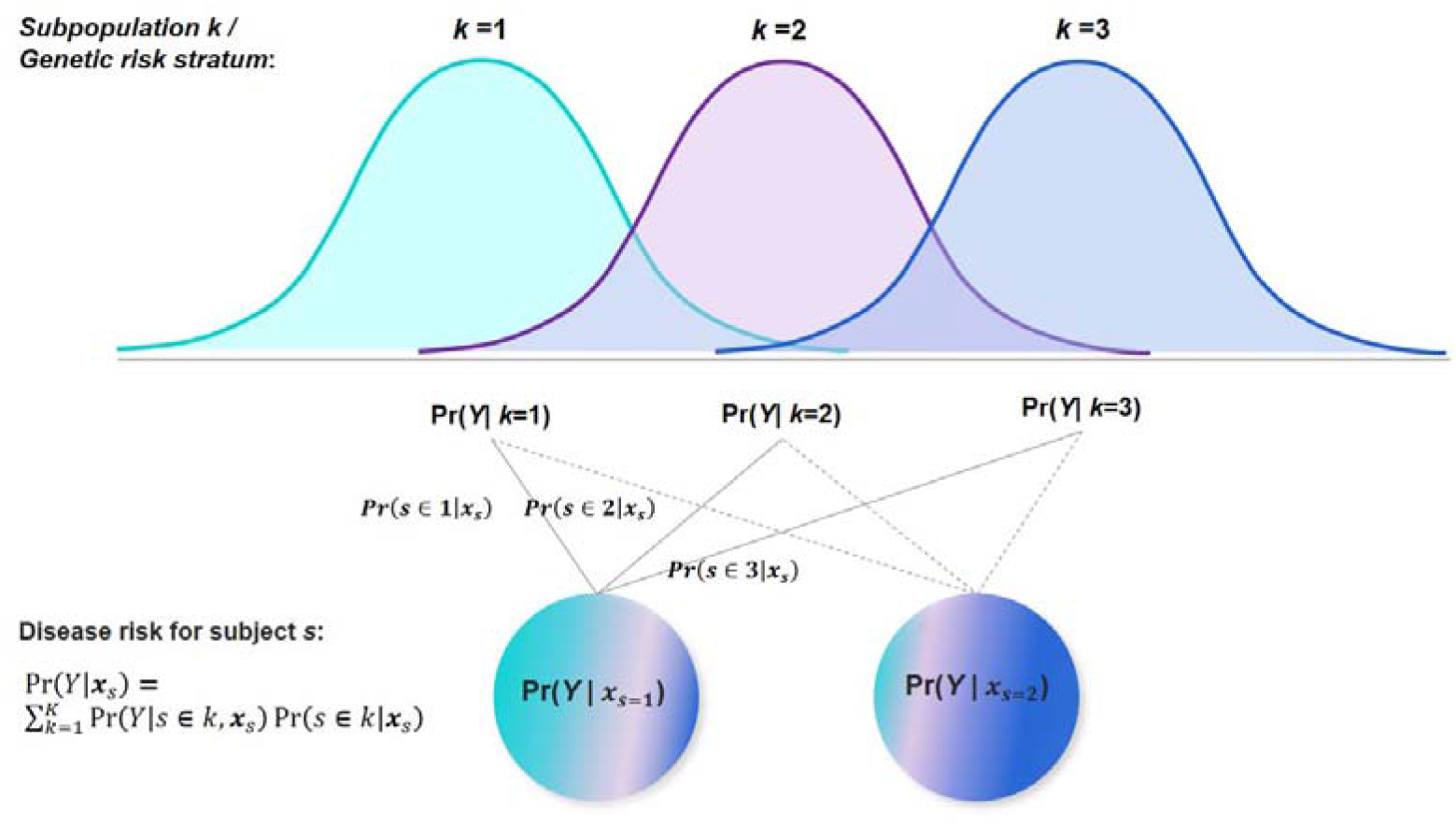
The Prism Vote (PV) framework for individualized risk prediction of traits. **Legend**: The PV views the complex trait of an individual as a composite risk outcome shaded from subpopulation strata, in which stratum-specific risk predictors may be estimated in a subpopulations by Pr (*Y*|*k*,***x***_*s*_), which are multiplied to a subject’s subpopulation propensity subpopulation comparatively more homogeneous. The stratum-wise risk is obtained in Pr (*s* ∈ *k*|***x***_*s*_). The total predicted risk of trait for an individual is the aggregated risk estimate from all subpopulations. In the PV framework, the dimension of individual identity is introduced. Each subject’s unique spectrum of propensities to subpopulations provides an individualized risk assessment outcome.

### PC-based population stratification and membership estimation

A PC-based approach is adopted to cluster subpopulations for its assumption-less property and adaptive feature in various data. Let *g*_*ij*_ be the genotype of subject *i* for SNP *j*, coded by minor allele counts (0, 1, 2), *I* = 1, ⋯, *N*, and *j* = 1, ⋯, *P*. The genetic matrix **G**_*N*×*P*_ is normalized to ***X***, by letting 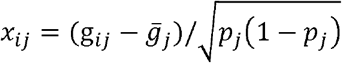, where 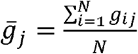 is the column mean of the SNP vector *j*, and 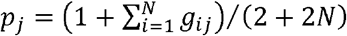 is the estimate of underlying allele frequency of SNP *j* (11). Compute the *N* × *N* covariance matrix **ψ**. Principal component analysis on ***X*** of both the training and testing data are performed for the individuals to obtain the eigenvectors, denoted as *ν^r^*, *r* = 1, ⋯, *N*, and **ψ*ν***_*r*_ = *λ*_*r*_***ν***_*r*_. Each vector *ν^r^* ∈ ℝ^*N*^ corresponds to the *r*^*th*^ largest eigenvalue 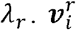 is the *i*^*th*^ loading of the eigenvector and carries the interpretation of *i*^*th*^ subject’s variation along the *r*^*th*^ ancestry axis. Suppose the top *q* eigenvectors contain a good amount of variation; 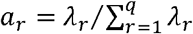 is the normalized eigenvalues. Compute the weighted score, 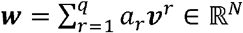,of which component *w*_*i*_ indicates the *i*^*th*^ individual’s variation summarized in the top *q* eigenvectors’ (ancestral) directions. By dividing ***w*** into *K* quantiles, the training subjects can be assigned to the *K* strata according to their quantile location in *w*. The eigenvectors were obtained using the PLINK (12).

The probability of observing a subject belonging to a particular stratum can be approximated based on the distance of a subject to the stratum center. The center of stratum *k* is 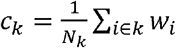, where *N*_*k*_ is the number of subjects in stratum *k*. Position of a new subject *s* in the ancestral space can be calculated from 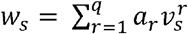. As the squared distance of a subject to a cluster center empirically follows a chi-squared distribution, the probability that subject *s* belongs to stratum *k* is estimated by,

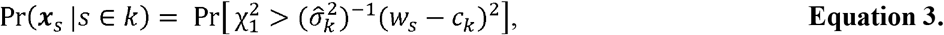

where 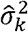 is the sample variance of *w*_*i*_ in the *k*^*th*^ stratum, *i* ∈ *k*. In the simulation and real data analysis, we set *q*=10 and *K* = 2.

In sum, the procedure of applying the PV is as follows: (1) Obtain eigenvectors and eigenvalues of all individuals genotypes (training and testing data) calculated in the ancestral direction; (2) divide the training set into *K* strata; (3) obtain stratum-wise predictors by a base prediction model in the training data, resulting *K* sets of predicted *Y* for the test data; (4) calculate the propensity of a test subject *s* to stratum *k* by **Equation 3**; (5) obtain the final predicted *Y* for the test set by **Equation 1**.

### Simulations

Simulation study I aims to investigate the effect of incorporating the PV with an LMM base prediction model in data consists of multiple populations. The genotype data was obtained from two real GWAS of the African and European populations from the GAIN project (dbGaP accession number: phs000021.v2.p1). Data including 1,932 African ancestry (AA) and 2,657 European ancestry (EA) subjects and 9,242 common (MAF>1%) genetic variants of chromosome 22 were extracted. An admixed population genotype data was simulated by sampling a genetic variant *x*_*i*, *j*_, for subject *i* at locus *j* from a binomial distribution, 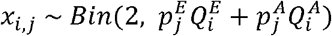, where 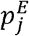 and 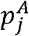 are MAF of locus *j* in the EA and AA data, respectively; 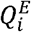 and 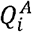 are the ancestry fractions of subject *i* in the two populations; 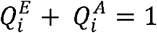 (**Supplementary S1**). Three thousand causal variants were selected for the AA and EA population, among which 75% was common to both populations and 25% was unique to a single population (13). Effect size was sampled from normal distributions with the number of variants in each effect group proportional to the effect magnitude. Specifically, the phenotype was determined by 10 SNPs of large effect from distribution *β* ~ N (0, 10^−2^); 300 SNPs of moderate effects from *β* ~ N (0, 10^−3^); and the remaining variants from *β* ~ N (0, 10^−4^) (14). Risk effect of the admixed population was simulated by setting 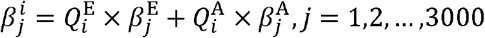, where 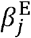 and 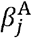 represent the effect sizes of SNP *j* in EA and AA, respectively. A linear model was used to obtain subjects’ phenotype from the set of causal variants and effect sizes, in which the residual term follows a normal distribution of variance satisfying alternative heritability scenarios (*h*^2^ = 0.2, 0.5 and 0.8). Simulation study II evaluates the performance of prediction algorithms as the similarity of effect size across population decreases. The genotype data was generated as described previously with 2,000 subjects from each of the EA, AA and admixed population, respectively. Genetic similarity was controlled by the covariance of the effect size in the EA and AA populations. Let ***β**^k^* ∈ ℝ^*m*^ denote the effects of *m* causal SNPs in population *k*. The coefficients follow a multivariate normal distribution (15),

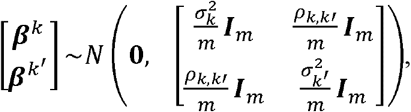

in which 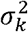 and 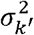, are the variance of the total additive genetic effect of these SNPs in two populations *k* and *k*′, *k* ≠ *k*′ ∈ {1, ⋯ *K*}. The covariance *ρ*_*k*,*k*′_ approximates the “shared heritability”. Thus, genetic similarity of two populations can be measured by the ratio *ρ*_*k*,*k*′_ /(*σ*_*k*_*σ*_*k*′_). When *η* = 0, the populations share no effect similarity; and as *η* approaches one, the traits are influenced by similar genetic effects in the mixed populations. Simulation study III considers sample size’s influence on prediction accuracy. In each heritability (*h*^2^ = 0.2, 0.5 and 0.8) by genetic similarity (*η* = 0.1, 0.5, 0.9) combination, eight datasets were simulated at different sample sizes (N = 1,000 to 40,000). Prediction accuracy is measured by the Pearson correlation coefficient between the observed and predicted outcome for continuous phenotypes, and by the area-under-the-curve (AUC) for binary outcomes. The mixed population data was divided into training and test sets in ratio of 4:1, and the average prediction accuracy calculated in five-group cross validations (5GCVs) was reported.

### Real data applications

The first GWAS data is from the Population Architecture through Genomics and Environment (PAGE) project (dbGap accession number phs000356.v1.p1). A total number 9,075 subjects were extracted, consisted of 3,520 self-identified African, 2,104 Hawaiians, and 3451 Japanese (**Supplementary S2**). Quality control (QC) was performed by removing SNPs with genotype call rate < 95%, Hardy-Weinberg equilibrium (HWE) *p*-value < 5 × 10^−8^, or MAF <0.01. After QC, 560,899 autosomal SNPs were available for analysis. The second GWAS is a subset of the non-European population in the UK Biobank (16). We included 5,718 individuals of Indian ancestry, 4,297 Caribbean, 3,204 African, 1,748 Pakistan 1,504 Chinese, 221 Bangladeshi, 2,869 admixed populations, and 6,947 subjects without clear ancestry information (**Supplementary S3, S4**). After the QC, 26,506 subjects and 524,557 SNPs were available for prediction study. In the PAGE data, traits including the body mass index (BMI), height, type II diabetes (diabetes) and hypertension were analyzed; and in the UK biobank data, BMI, diabetes diagnosed by doctor (diabetes), and the cardiovascular disease (CVD) outcomes were analyzed.

## Results

### Simulation Study I: the PV using linear mixed model as prediction base

In this simulation, prediction accuracy was evaluated in mix population cohorts by the PV, implemented with alternative base prediction models (base model), and by the reference methods using the base models controlling for the top 10 PCs. For the base models, we consider the linear regression model (LM), the BayesR (14), and the Dirichlet Process Regression (DPR) (17). The BayesR assumes that the variant effect follows a normal mixture distribution with the majority of variants having no effect on the phenotype. On the contrary, the DPR adopts a non-parametric prior on the effect distributions and assigns non-zero effect on all variants. In the mixed population, true ancestry information was unknown and was controlled purely through statistical modelling. **Figure 2** showed that the PV generally improved prediction accuracy of the base models comparing to the reference group in 5GCV. Under the high heritability scenario, the PV improved the mean prediction correlation coefficient of the BayesR from 0.49 to 0.54 by 10.2%, and improved the DPR from 0.46 to 0.58 by 26.5% (**Supplementary S5**). The PV improved the LM and DPR in all heritability scenarios, while it only enhanced the BayesR under the high heritability setting. This might be due to the sparse effect model assumption made by the BayesR, from which modest causal effects were prevailingly estimated as zero in the low heritability data.

**Figure 2.**
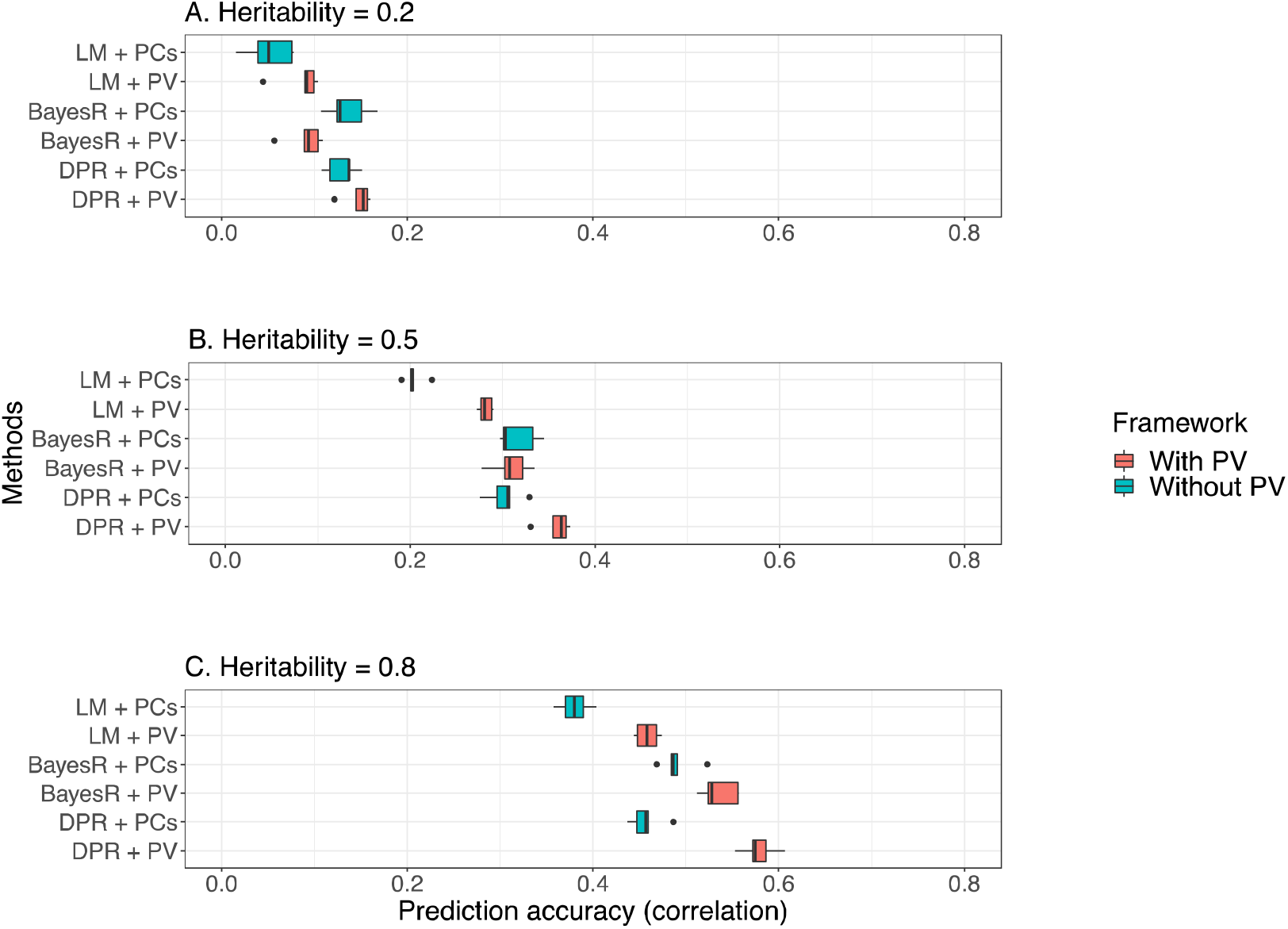
Prediction outcome of the Prism Vote implemented with alternative base prediction models (Simulation Study I) **Legend**: Panel (A): Heritability = 0.2; (B): Heritability = 0.5; (C): Heritability = 0.8. Prediction performance of the PV with the base models are compared to reference methods. The three base models implemented are the linear regression model (LM), BayesR, and the Dirichlet process regression (DPR). In all scenarios, the PV exhibited higher mean prediction accuracy in terms of concordance correlation coefficient in 5GCV compared to the reference, and PV’s advantage enlarged as the heritability in the data increased. **Supplementary S6** includes prediction outcomes in single populations.

### Simulation Study II and III: Prediction as genetic heterogeneity and sample size increased

Next, we investigate the performance of the PV with the DPR base as genetic heterogeneity across populations varied. As shown in **Figure 3**, as the genetic similarity in populations decreased, the reference group (DPR + PCs)’s prediction accuracy observed a significant reduction, while the DPR implemented in the PV framework produced a stable prediction accuracy in all scenarios. For instance, under the high heritability scenario (**Figure 3C**), as the effect similarity decreased from 80% to 20%, the mean prediction accuracy by the base model in the reference group reduced from 0.52 to 0.40 by 23.1%, while the accuracy with PV only slightly dropped from 0.48 to 0.47 by 2.1%. In Figure 3B and 3C’s medium and high heritability settings, prediction gain by the PV was warranted when the genetic similarity in multiple populations was lower than half. In simulation III (**Figure 4**), sample size of data steadily increased from 1,000 to 40,000 in different combinations of heritability (*h*^2^ = 0.2, 0.5 and 0.8) and effect similarity (*η* = 0.1, 0.5, 0.9) levels. In large samples, implementing the PV resulted in prediction accuracy gain in all nine scenarios (**Figure 4** and **Supplementary materials S7-8**). Particularly, when *N*=40,000, *h*^2^= 0.8 and η = 0.1, the PV improved the prediction accuracy of the DPR from 0.59 to 0.80 by 26.3%.

**Figure 3.**
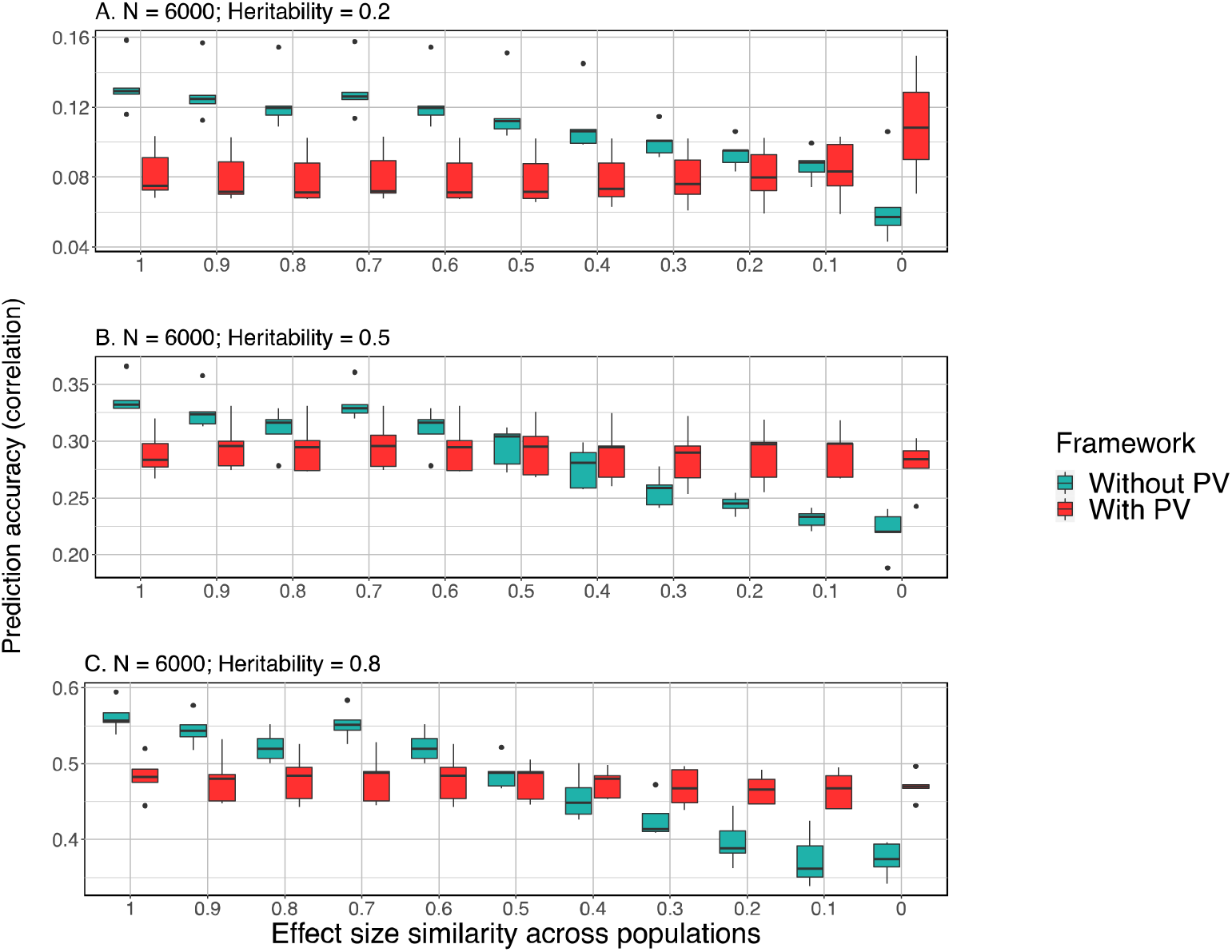
Prediction performance of PV as genetic heterogeneity increases across populations (Simulation Study II) **Legend**: With PV: Prediction model is the Prism Vote with the DPR base model; Without PV (the reference method): DPR controlling for the top 10 PCs. Panel (A): Heritability = 0.2; (B): Heritability = 0.5; (C): Heritability = 0.8. Vertical axis: average correlation coefficient of the predicted and observed phenotype in 5GCVs. Horizontal axis: levels of effect size similarity across populations. As effect size similarity decreased across populations, the PV showed stable prediction outcomes (red), while performances of the reference method were affected substantially (green).

**Figure 4.**
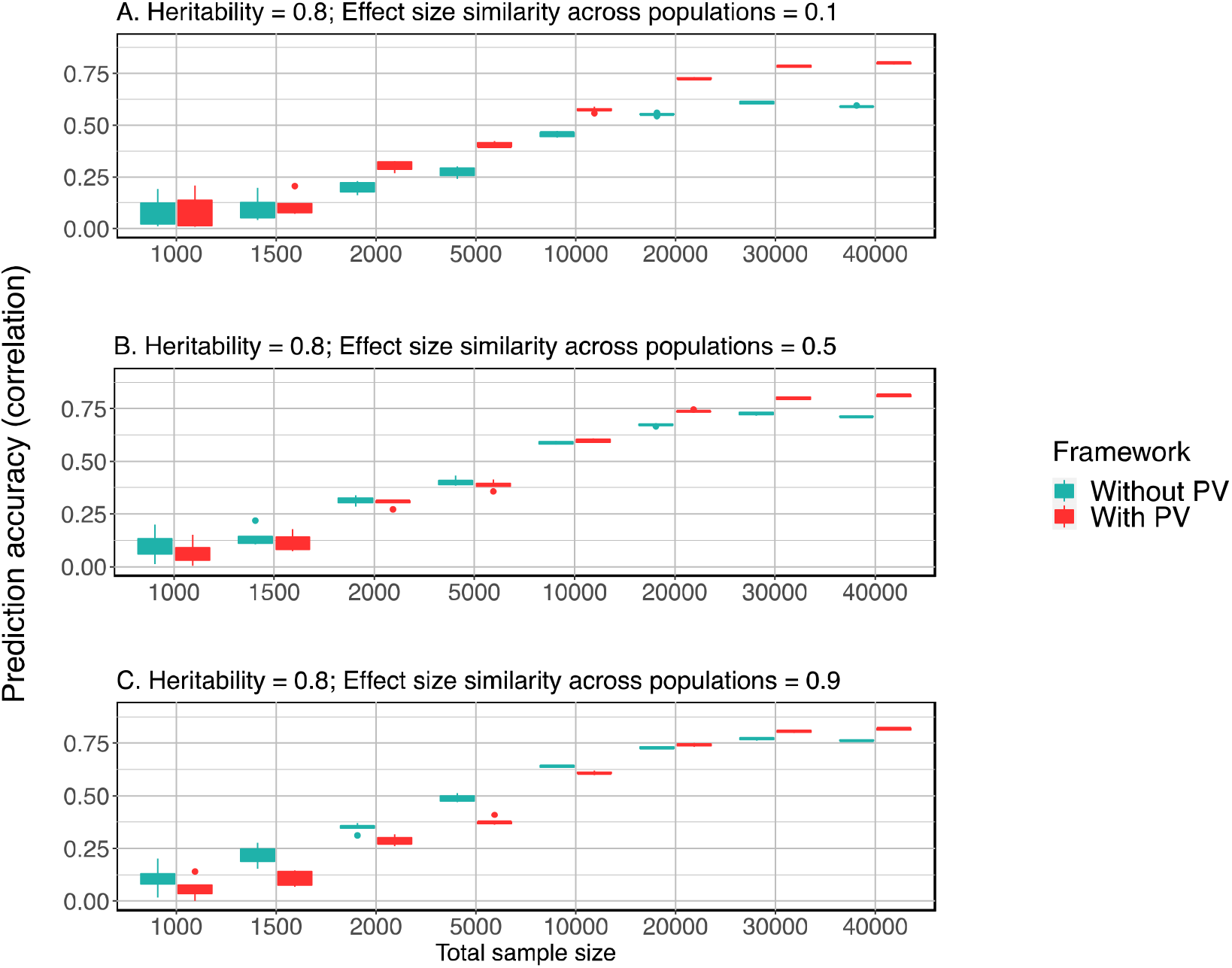
Prediction performance of PV as sample size changes (Simulation study III) **Legend**: Prediction performance of PV and the reference method using the DPR base is evaluated at different sample sizes, heritability and genetic heterogeneity levels. Panel (A): Heritability = 0.8, effect size similarity *η* = 0.1; (B): Heritability = 0.8, *η* = 0.5; (C): Heritability = 0.8, *η* = 0.9. As the sample size increased from 1,000 to 40,000, the prediction accuracy of PV continued to rise. The PV’s advantage was more obvious when the genetic heterogeneity was high (Panel A and B). Results for scenarios of heritability = 0.2 and 0.5 can be found in **Supplementary materials S7-8**.

### Prediction in Real GWAS Datasets

In the first real GWAS data from the PAGE project, the PV was implemented with the DPR for predicting BMI, height, diabetes and hypertension (**Figure 5**). Comparing to the reference method of DPR and PCs, the PV enhanced the prediction accuracy by 12.1% (SD 4.7%) for the BMI, 2.0% (1.5%) for height, 5.2% (SD 2.1%) for hypertension, and 5.4% (SD 2.2%) for diabetes. For easier interpretation of the results, we also displayed the prediction outcome achieved in single populations (**Figure 5**). In general, by the reference method, the prediction accuracy in the joint cohort was in between the highest and lowest performance achieved in a single population (**Figure 5B-D**), while the PV elevated the prediction outcome in joint data close to the best accuracy reached in a single population. For example, in **Figure 5D**, the prediction for diabetes by the reference method was in between its performance in the Japanese population observing the lowest accuracy and the African population the second lowest; while the PV improved the joint cohort prediction to an accuracy achieved in the Hawaiian, the population observing the highest performance. Furthermore, as shown in **Figure 5A**, the PV increased the prediction accuracy for the BMI to 0.374 (SD 0.004) in the mixed population, a level unreached in single populations, among which the best performance was only 0.214 (SD 0.136). In the UK Biobank data composed of five minority populations, applying the PV with the DPR base significantly improved the prediction accuracy for the BMI by 12.0% (SD 5.6%), for the CVD by 5.9% (SD 1.0%), and for the diabetes by 3.7% (SD 2.2%) (**Figure 6**). Prediction standard deviations also considerably reduced in the joint data analysis benefited from the larger sample size. Finally, we compared the estimated effect size of the top 5,000 SNPs in the two real GWAS datasets (**Supplementary materials S11-12**); the effect sizes were vastly different between strata, suggesting prevailing genetic heterogeneity in these data.

**Figure 5.**
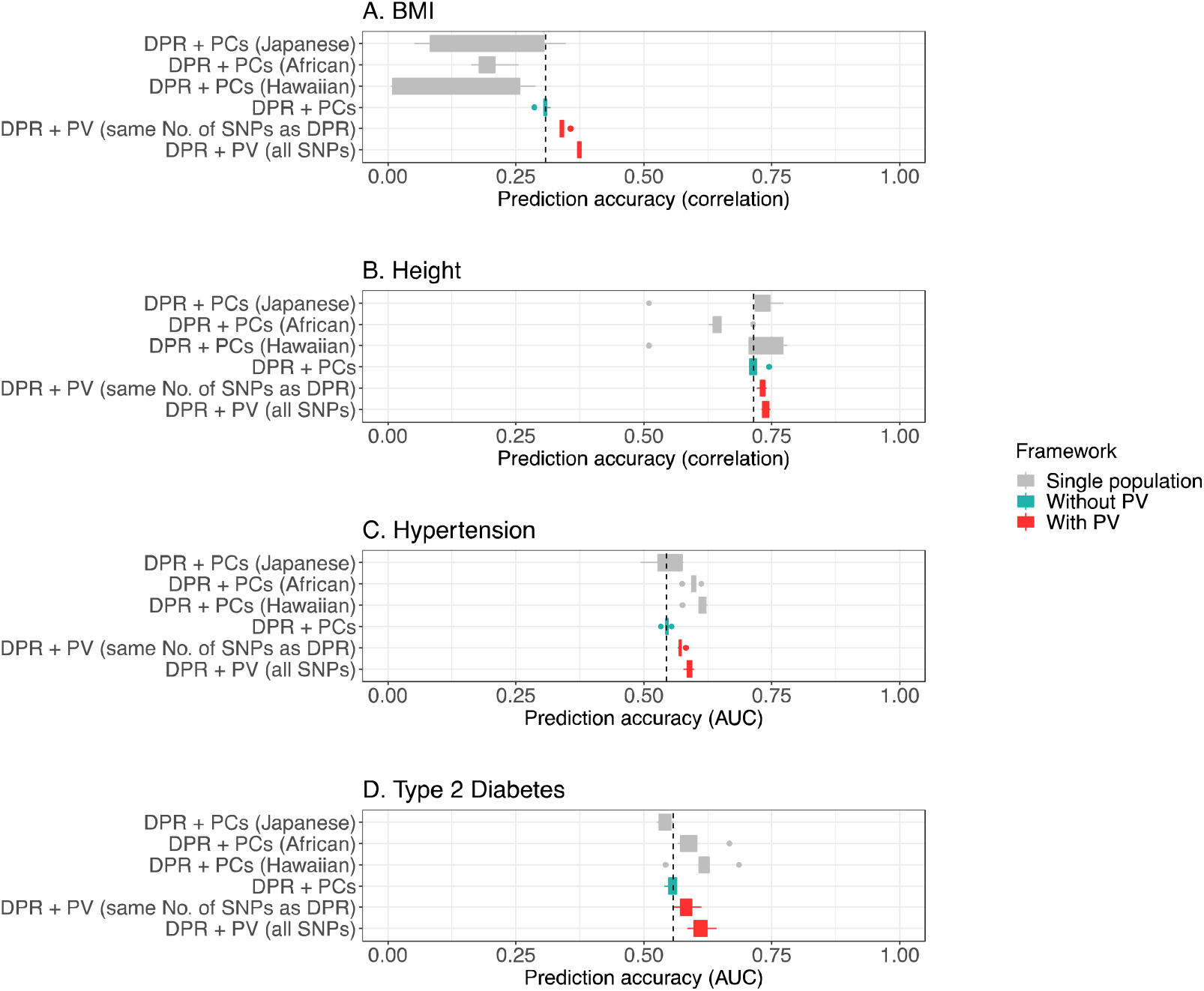
Prediction performance of the PV in real data application (PAGE dataset) **Legend:** Mean 5GCV Prediction accuracy of the PV and reference method using the DPR base model in the PAGE mixed population data. Comparing to the reference, the PV enhanced the prediction accuracy of DPR by 12.1% (SD 4.7%) for the BMI, 2.0% (1.5%) for height, 5.2% (SD 2.1%) for hypertension, and 5.4% (SD 2.2%) for diabetes. Prediction outcome in single populations by the reference method is shown in gray color. Details can be found in **Supplementary material S9**.

**Figure 6.**
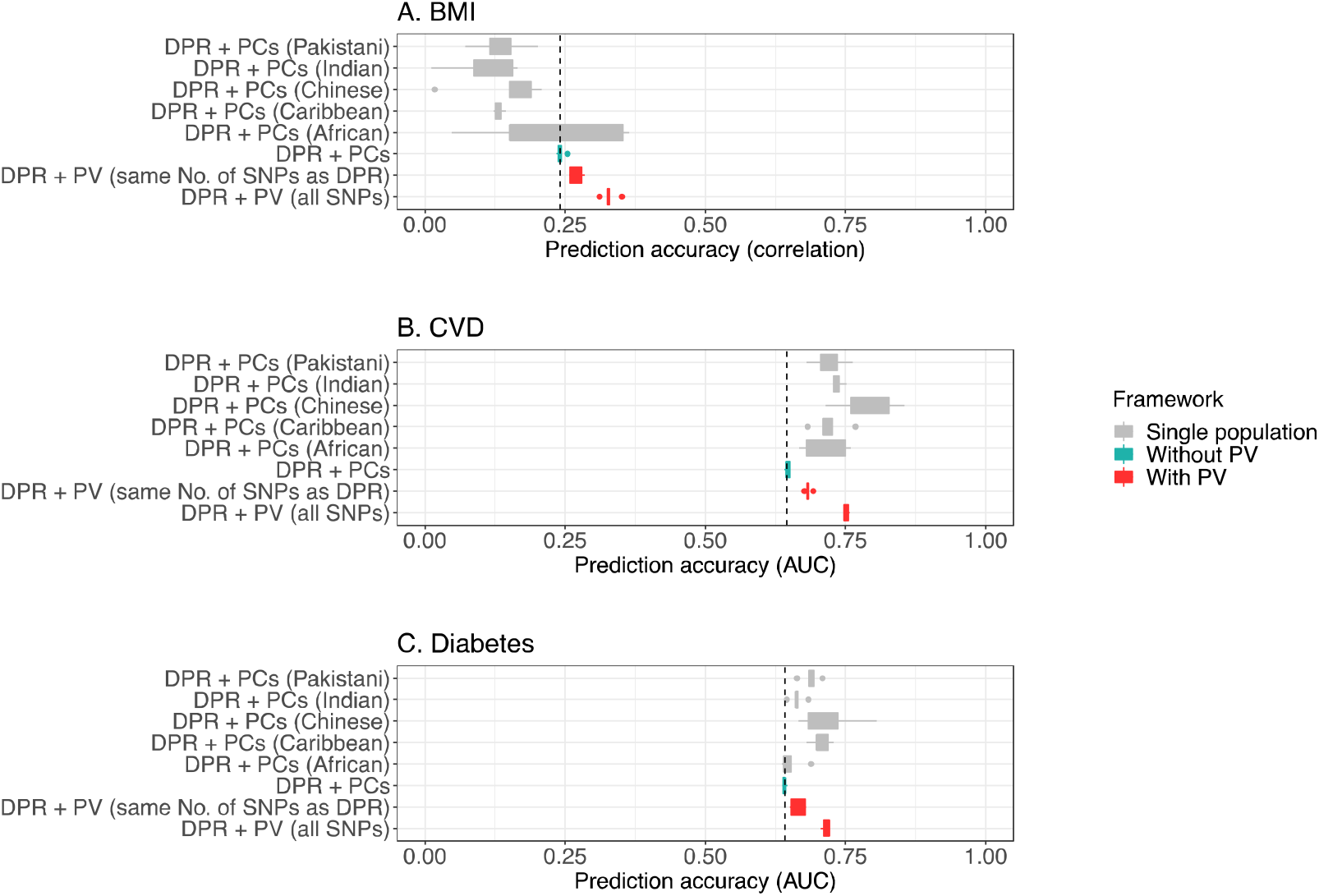
Prediction performance of the PV in real data application (UK Biobank data) **Legend**: Mean 5GCV prediction accuracy of the PV and reference method using the DPR base model in the UK Biobank data. Applying the PV significantly improved the prediction accuracy of the DPR for the BMI by 12.0% (SD 5.6%), for the CVD by 5.9% (SD 1.0%), and for the diabetes by 3.7% (SD 2.2%). Details can be found in **Supplementary material S10**.

## Discussion

In this study, the Prism Vote framework is introduced to dissect and integrate the risk of individuals based on personalized risk spectrum through a Bayesian probability framework. Simulation studies showed that the method generally enhanced the prediction of base models in different heritability scenarios, and advantage of the framework expanded with increasing genetic heterogeneity and sample size. Application of the PV in two real mixed population GWASs also resulted significant gain in prediction accuracy.

The component steps of the framework can be substituted with alternative methods according to data attributes. In the population stratification step, either a model-based or model-free model may be incorporated (11, 18, 19). The two approaches were tested on a simulated admixed cohort generated from two distinct populations (**Supplementary Notes**, **Supplementary material S13**). Individuals’ group membership probabilities obtained by the PC-based method described in this study gave concordant estimates as the outcome obtained from the Bayesian maximum likelihood approach implemented in the ADMIXTURE software (19) (**Supplementary material S14**). For the prediction step, the linear mixed model was suitable for this study design for its good property of simultaneous estimation of whole genome SNPs effects and prediction. Other approaches, such as the machine learning methods, or the PRS, may be applied to construct predictors in stratum. To apply the PRS, the following issues shall be considered. The PRS draws summary statistics from well-powered external datasets, however, these external populations were predominantly of the European ancestries. Therefore, although coefficient of the aggregated PRS score can be evaluated in stratum, to compute the differentiation of SNP-effects across stratum, one may need to estimate the transferrable part of the effect size from auxiliary data to strata (10, 20). Alternatively, the stratum specific SNP-effects can always be estimated directly from applying a LMM in stratum, followed by constructing the stratum’s PRS.

The PV framework leverages on the data’s genetic architecture to form homogeneous genetic strata. The grouping of subjects and the optimal number of strata is a complicated issue, as it is simultaneously influenced by the sample size, underlying genetic models of the trait, and genetic architecture in strata. In either the model-based or model-free approach, the number of population clusters was often determined empirically (5, 11, 18). In the current analysis, an equal-division approached was adopted such that each stratum has the same sample size. However, the division step could be further optimized considering the bias-variance tradeoff for within stratum SNP-effects estimation towards achieving optimal prediction outcome, which requires extensive research in future study.

Another potential advantage of the PV is that it allows distributed computing of large genomic datasets in the dimension of subjects. Traditionally, SNPs are assigned to multiple clusters to increase calculation efficiency, however, parallel computing is difficult to realize for models performing simultaneous evaluation of SNPs in a single model, such as the LMM or Lasso. However, the PV framework, instead of distributing the biomarkers, may assign subgroups of subjects to CPU-clusters and at the same time maintaining the total information gain in the final prediction outcome. Finally, the framework may also be extended to consider the subgrouping based on related traits’ genetic structure to leverage on the pleiotropy effect of genes for multiple phenotypes.

## Disclaimer

The funding agencies had no role in the design and conduct of the study; collection, management, analysis, and interpretation of the data; preparation, review, or approval of the manuscript; or decision to submit the manuscript for publication.

## Authors’ contributions

MHW and XX conceived the study. XX carried out the analysis. QL, RS, MKCC, WKKW, YW, HT, BCYZ discussed the results. HT helped design the simulation study. MHW and XX drafted the first manuscript. All authors critically read and revised the manuscript and gave final approval for publication.

## Acknowledgments

**The UK Biobank data:** We conducted the research using the UKBB resource under approved data requests (refs: 57883).

**The Population Architecture through Genomics and Environment (PAGE) data**: Funding support for "Epidemiologic Architecture for Genes Linked to Environment (EAGLE)" was provided through the National Human Genome Research Institute's Population Architecture Using Genomics and Epidemiology (PAGE) network (U01HG004798-01). The human subjects participating in the study derive from the National Health and Nutrition Examination Surveys, and these studies are supported by the Centers for Disease Control and Prevention. Funding support for the PAGE Multiethnic Cohort study was provided through the National Cancer Institute (R37CA54281, R01 CA63, P01CA33619, U01CA136792, and U01CA98758) and the National Human Genome Research Institute (U01HG004802). Funding support for the “Epidemiology of putative genetic variants: The Women’s Health Initiative” was provided through the National Human Genome Research Institute's Population Architecture Using Genomics and Epidemiology (PAGE) network (U01HG004790). The WHI program is funded by the National Heart, Lung, and Blood Institute; NIH; and U.S. Department of Health and Human Services through contracts N01WH22110, 24152, 32100-2, 32105-6, 32108-9, 32111-13, 32115, 32118-32119, 32122, 42107-26, 42129-32, and 44221. Funding support for the Genetic Epidemiology of Causal Variants Across the Life Course (CALiCo) was provided through the National Human Genome Research Institute's Population Architecture Using Genomics and Epidemiology (PAGE) network (U01HG004803). The human subjects derive from the following studies: Atherosclerosis Risk in Communities (ARIC) Study, Coronary Artery Risk Development in Young Adults (CARDIA), and Cardiovascular Health Study (CHS). The Atherosclerosis Risk in Communities (ARIC) Study is carried out as a collaborative study supported by National Heart, Lung, and Blood Institute contracts N01-HC-55015, N01-HC-55016, N01-HC-55018, N01-HC-55019, N01-HC-55020, N01-HC-55021, N01-HC-55022. The Coronary Artery Risk Development I Young Adults (CARDIA) study is supported by the following National Institutes of Health, National Heart, Lung and Blood Institute contracts: N01-HC-95095; N01-HC-48047; N01-HC-48048; N01-HC-48049; N01-HC-48050; N01-HC-45134; N01-HC-05187; and N01-HC-45205. The Cardiovascular Health Study (CHS) is supported by contracts N01-HC-35129, N01-HC-45133, N01-HC-75150, N01-HC-85079 through N01-HC-85086, N01 HC-15103, N01 HC-55222, and U01 HL080295 from the National Heart, Lung, and Blood Institute, with additional contribution from the National Institute of Neurological Disorders and Stroke and grant AG09556 from the National Institute of Aging Assistance with phenotype harmonization, SNP selection, data cleaning, meta-analyses, data management and dissemination, and general study coordination, was provided by the PAGE Coordinating Center (U01HG004801-01). The datasets used for the analyses described in this manuscript were obtained from dbGaP at phs000356.v1.p1.

**The Genetic Association Information Network Schizophrenia data:** Funding support for the Genome-Wide Association of Schizophrenia Study was provided by the National Institute of Mental Health (R01 MH67257, R01 MH59588, R01 MH59571, R01 MH59565, R01 MH59587, R01 MH60870, R01 MH59566, R01 MH59586, R01 MH61675, R01 MH60879, R01 MH81800, U01 MH46276, U01 MH46289 U01 MH46318, U01 MH79469, and U01 MH79470) and the genotyping of samples was provided through the Genetic Association Information Network (GAIN). The datasets used for the analyses described in this manuscript were obtained from the database of Genotypes and Phenotypes (dbGaP) found at http://www.ncbi.nlm.nih.gov/gap through dbGaP accession number phs000021.v2.p1. Samples and associated phenotype data for the Genome-Wide Association of Schizophrenia Study were provided by the Molecular Genetics of Schizophrenia Collaboration (PI: Pablo V. Gejman, Evanston Northwestern Healthcare (ENH) and Northwestern University, Evanston, IL, USA).

