## Supplementary Figures, Tables and Notes for "A Prism Vote Framework for Individualized Risk Prediction of Traits in Genome-wide Sequencing Data of Multiple Populations"

**Supplementary Materials**

**S1: The genetic ancestries in simulation study I**

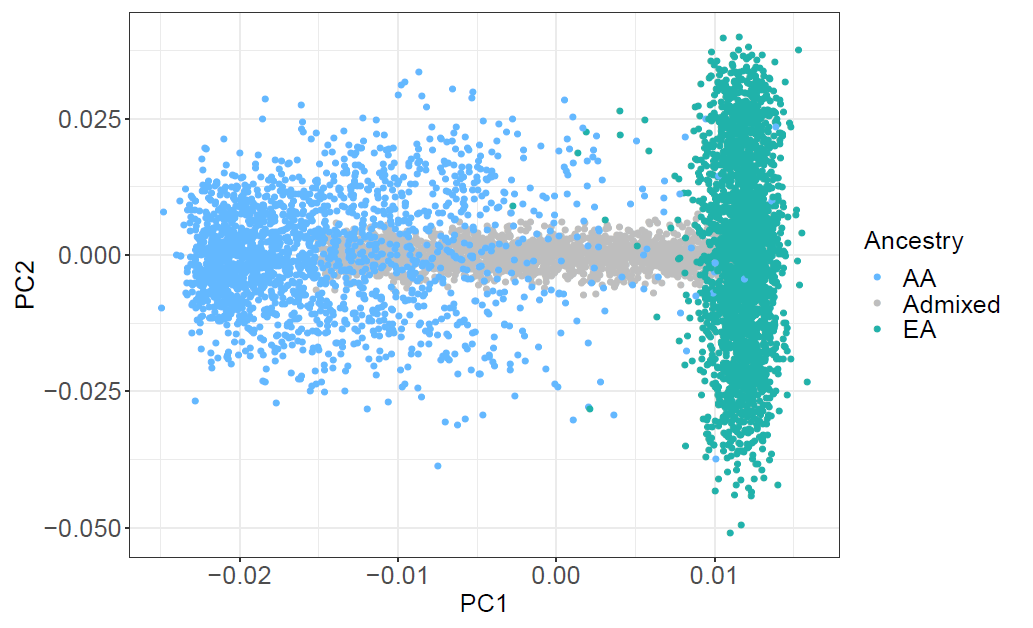

**Legend**: The three populations (European ancestry (EA), African ancestry (AA), Admixed population (Admixed)) plotted in the coordinates spanned by the top two principal components (PCs).

**S2.** **The genetic ancestries in PAGE data**

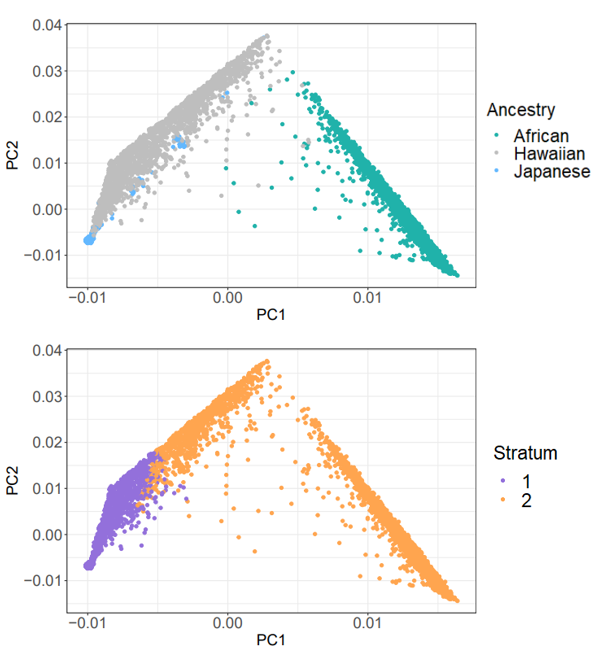

**Legend:** A. The populations plotted in the coordinates spanned by the top two principal components (PCs) annotated by ancestry information. B. The distribution of subjects in two strata when we stratify all subjects by the median of weighted sum of top 10 PCs.

**S3: The genetic ancestry of minority populations in UKbiobank.**

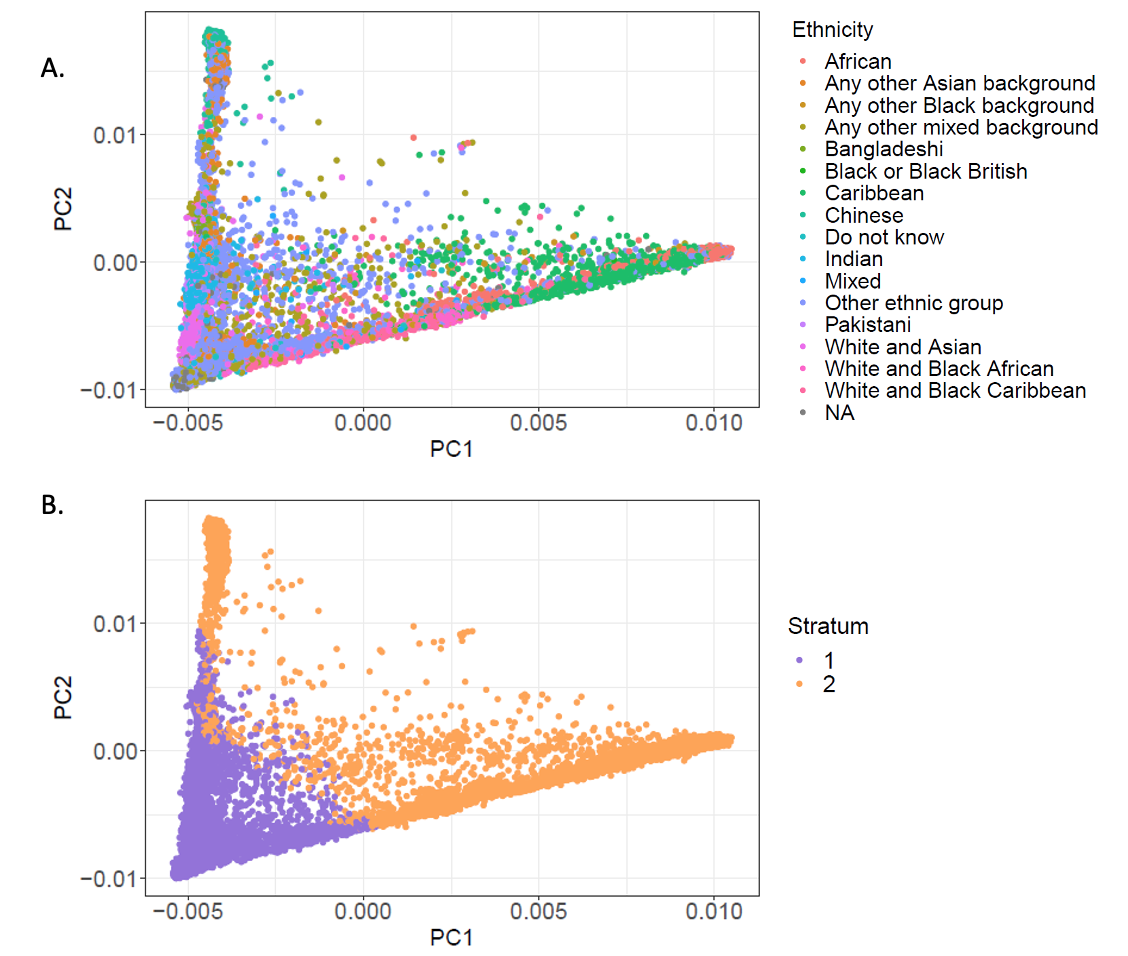

**Legend:** A. The populations plotted in the coordinates spanned by the top two principal components (PCs) annotated by ancestry information. B. The distribution of subjects in two strata when we stratify all subjects by the median of weighted sum of top 10 PCs.

**S4: The sample size in non-European populations in UK Biobank.**

| **Populations** | **Sample size** |
| --- | --- |
| Indian | 5716 |
| Other ethnic group | 4356 |
| Caribbean | 4297 |
| African | 3204 |
| Pakistani | 1748 |
| Any other Asian background | 1747 |
| Chinese | 1504 |
| Any other mixed background | 996 |
| White and Asian | 802 |
| White and Black Caribbean | 597 |
| NA | 522 |
| White and Black African | 402 |
| Bangladeshi | 221 |
| Do not know | 204 |
| Any other Black background | 118 |
| Mixed | 46 |
| Black or Black British | 26 |

**S5. The Pearson correlation of predicted phenotype and true phenotype using different methods (Simulation Study I).**

| Heritability | 0.2 | 0.5 | 0.8 |
| --- | --- | --- | --- |
| LM + PCs | 0.052 (0.026)^*^ | 0.204 (0.012) | 0.380 (0.018) |
| LM + PV | 0.086 (0.024) | 0.281 (0.008) | 0.458 (0.013) |
| BayesR + PCs | 0.136 (0.024) | 0.316 (0.022) | 0.491 (0.020) |
| BayesR + PV | 0.090 (0.020) | 0.309 (0.021) | 0.535 (0.020) |
| DPR + PCs | 0.130 (0.017) | 0.302 (0.020) | 0.458 (0.018) |
| DPR + PV | 0.147 (0.015) | 0.358 (0.017) | 0.579 (0.020) |

^*^: The standard deviation of prediction correlation was listed in brackets.

**S6. Compare combined analysis with separate analysis**

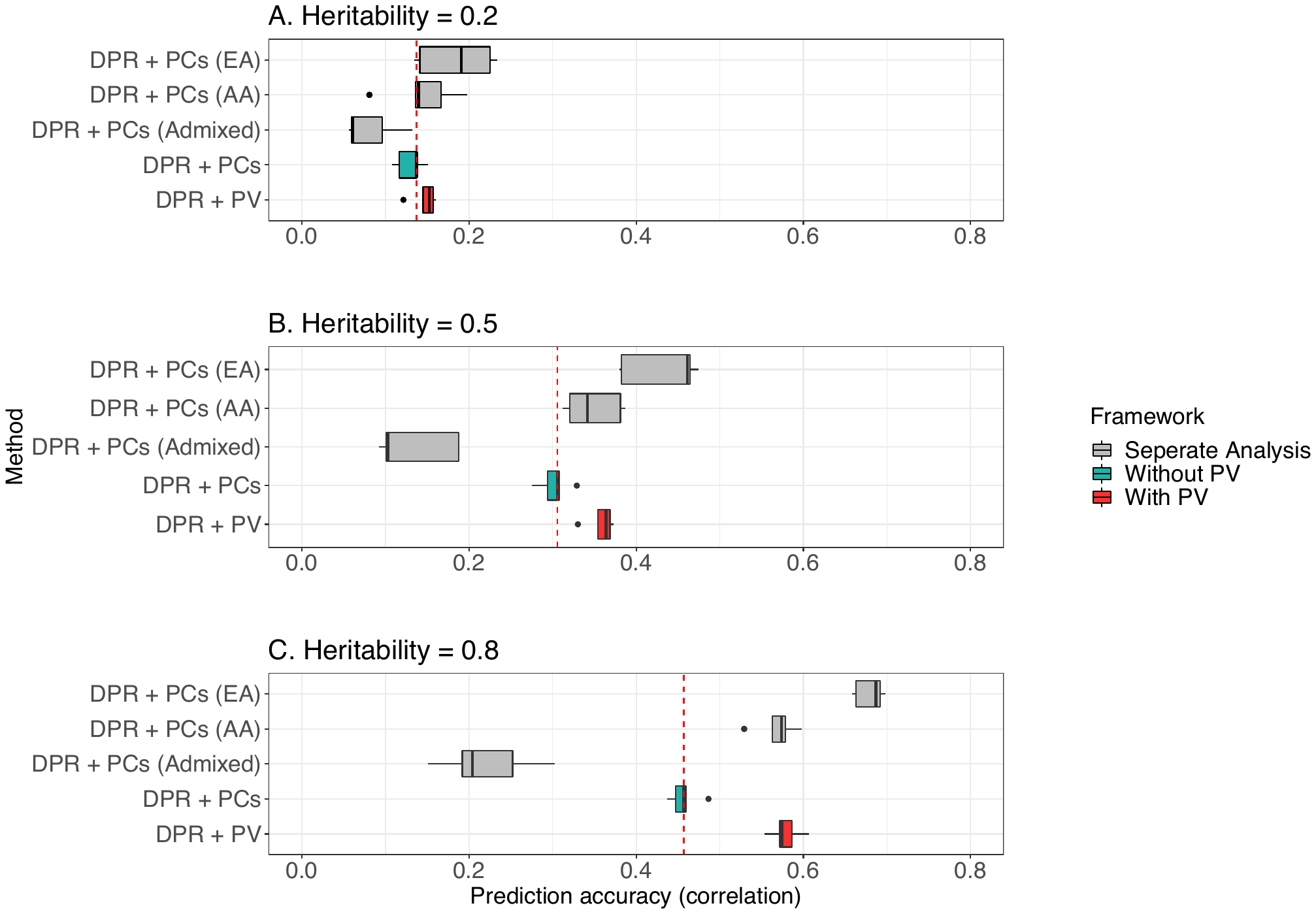

**Legend: Simulation study I:** Comparison of prediction performance of combined analysis with separate analysis in three simulated population (EA: European Ancestry, AA: African Ancestry, Admixed: admixed population). Performance is measured by Pearson correlation of predicted phenotype with real phenotype. Among three populations, admixed population benefits the most from combined analysis in all heritability settings. The prediction Pearson correlation in admixed population in three heritability settings are, respectively, 0.08(0.03), 0.13(0.05), 0.22(0.06), while combined analysis achieved 0.13(0.02), 0.30(0.02), 0.46(0.02) for DPR + PCs, and 0.15(0.02), 0.36(0.02), 0.58(0.02) for DPR + PV.

**S7. The prediction performance of PV with increasing sample size when heritability is 0.5.**

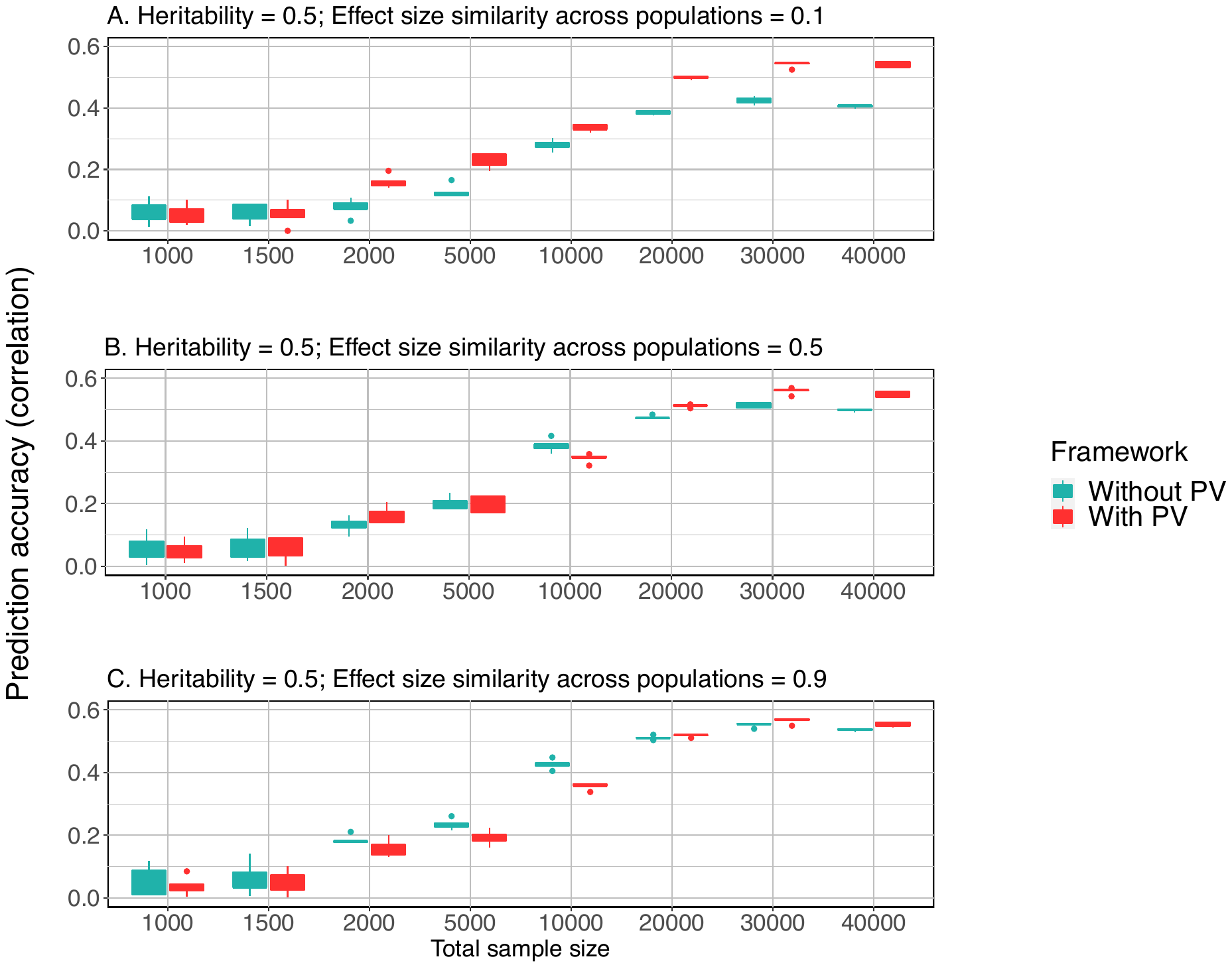

**Legend: Simulation study III:** The prediction performance of DPR + PV and DPR + PCs with increasing sample size when heritability = 0.5. Performance is measured by Pearson correlation of predicted phenotype with real phenotype. With increasing sample from 1,000 to 40,000, the improvement of DPR + PV over DPR + PCs are more obvious, even when the shared proportion of heritability = 90%.

**S8: The prediction performance of PV with increasing sample size when heritability is 0.2.**

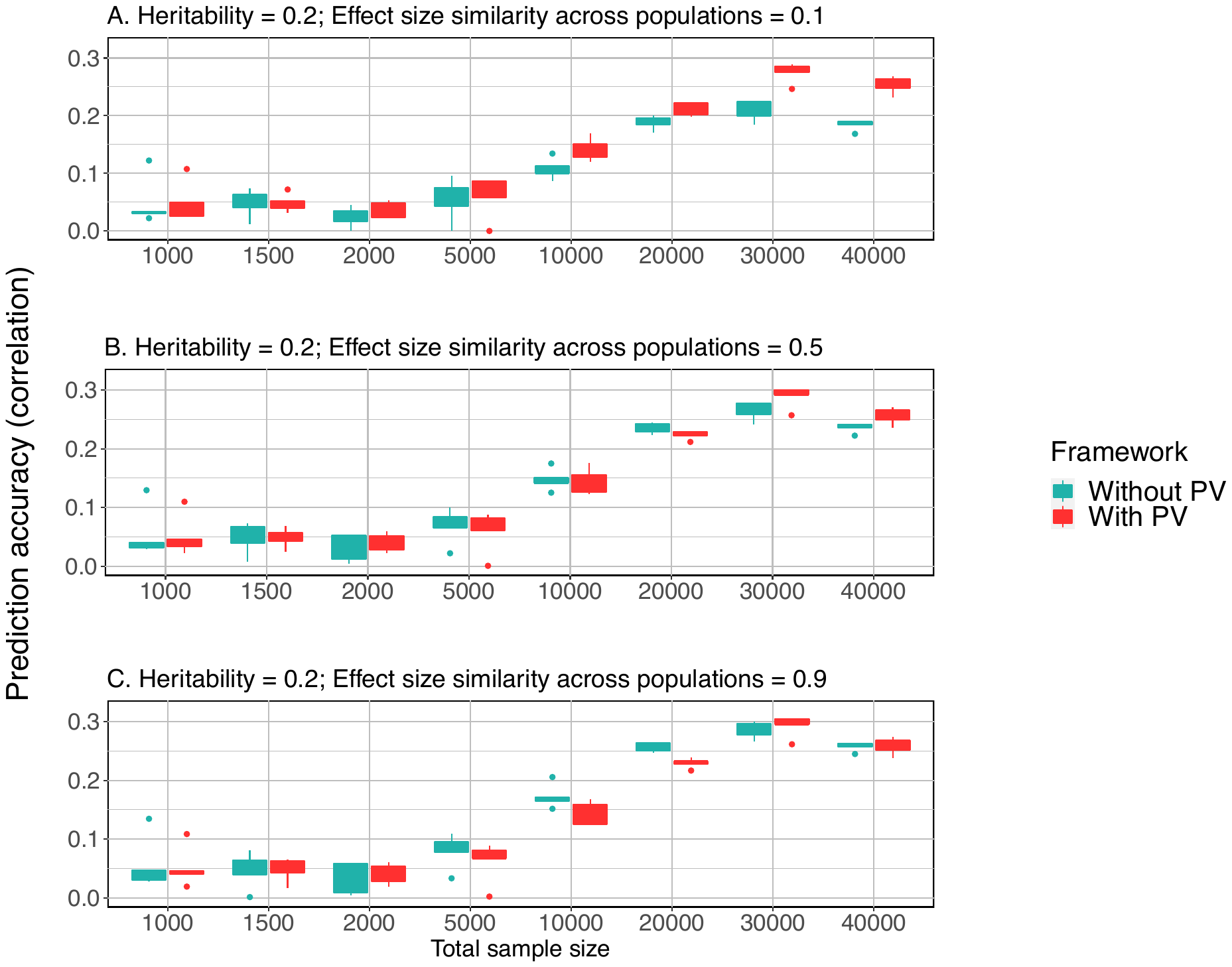

**Legend: Simulation study III:** The prediction performance of DPR + PV and DPR + PCs with increasing sample size when heritability = 0.2. Performance is measured by Pearson correlation of predicted phenotype with real phenotype. With increasing sample from 1,000 to 40,000, the improvement of PV over just controlling PCs are more obvious.

**S9. Prediction performance in PAGE data using different analysis framework.**

|  | BMI | Height | Hypertension | Type 2 Diabetes |
| --- | --- | --- | --- | --- |
| DPR + PCs^*^ | 0.305  (0.012)^+^ | 0.718  (0.016) | 0.544  (0.008) | 0.555  (0.011) |
| DPR + PV  (same No. of SNPs as DPR)^*^ | 0.342  (0.009) | 0.732  (0.007) | 0.572  (0.006) | 0.585  (0.022) |
| DPR + PV  (all SNPs) | 0.374  (0.004) | 0.739  (0.007) | 0.589  (0.008) | 0.612  (0.022) |
| DPR + PCs  (Hawaii, all SNPs) | 0.136  (0.133) | 0.704  (0.112) | 0.610  (0.020) | 0.616  (0.051) |
| DPR + PCs  (African, all SNPs) | 0.197  (0.036) | 0.652  (0.035) | 0.596  (0.014) | 0.600  (0.040) |
| DPR + PCs  (Japanese, all SNPs) | 0.214  (0.136) | 0.698  (0.107) | 0.549  (0.037) | 0.542  (0.014) |

***:** Due to computation burden, SNPs with p-value less than 0.1 were used in DPR + PCs; to achieve relatively fair comparison, when applying PV to DPR, we extracted the same number of SNPs in each stratum and marked as DPR + PV (same No. of SNPs as DPR).

**+:** Standard deviation of 5GCVs prediction performance.

The prediction accuracy measure is the Pearson correlation coefficient for BMI and Height, and the area-under-the-curve (AUC) statistic for Hypertension and Type 2 Diabetes.

**S10. Prediction performance in UK Biobank data using different analysis framework.**

|  | BMI | CVD | Diabetes |
| --- | --- | --- | --- |
| DPR + PCs^*^ | 0.242  (0.007)^+^ | 0.646  (0.004) | 0.641  (0.003) |
| DPR + PV  (same No. of SNPs as DPR)^*^ | 0.271  (0.013) | 0.684  (0.006) | 0.666  (0.014) |
| DPR + PV  (all SNPs) | 0.329  (0.014) | 0.751  (0.005) | 0.716  (0.007) |
| DPR + PCs  (African, all SNPs) | 0.215  (0.122) | 0.638  (0.031) | 0.610  (0.027) |
| DPR + PCs  (Caribbean, all SNPs) | 0.119  (0.021) | 0.670  (0.031) | 0.644  (0.018) |
| DPR + PCs  (Chinese, all SNPs) | 0.150  (0.078) | 0.772  (0.080) | 0.671  (0.060) |
| DPR + PCs  (Indian, all SNPs) | 0.111  (0.026) | 0.683  (0.009) | 0.638  (0.013) |
| DPR + PCs  (Pakistani, all SNPs) | 0.172  (0.056) | 0.706  (0.038) | 0.676  (0.027) |

***:** Due to computation burden, SNPs with p-value less than 0.1 were used in DPR + PCs; to achieve relatively fair comparison, when applying PV to DPR, we extracted the same number of SNPs in each stratum and marked as DPR + PV (same No. of SNPs as DPR).

**+:** Standard deviation of 5GCVs prediction performance.

The prediction accuracy measure is the Pearson correlation coefficient for BMI, and the area-under-the-curve (AUC) statistic for CVD and Diabetes.

**S11. Effect size stratification by subpopulations in PAGE data.**

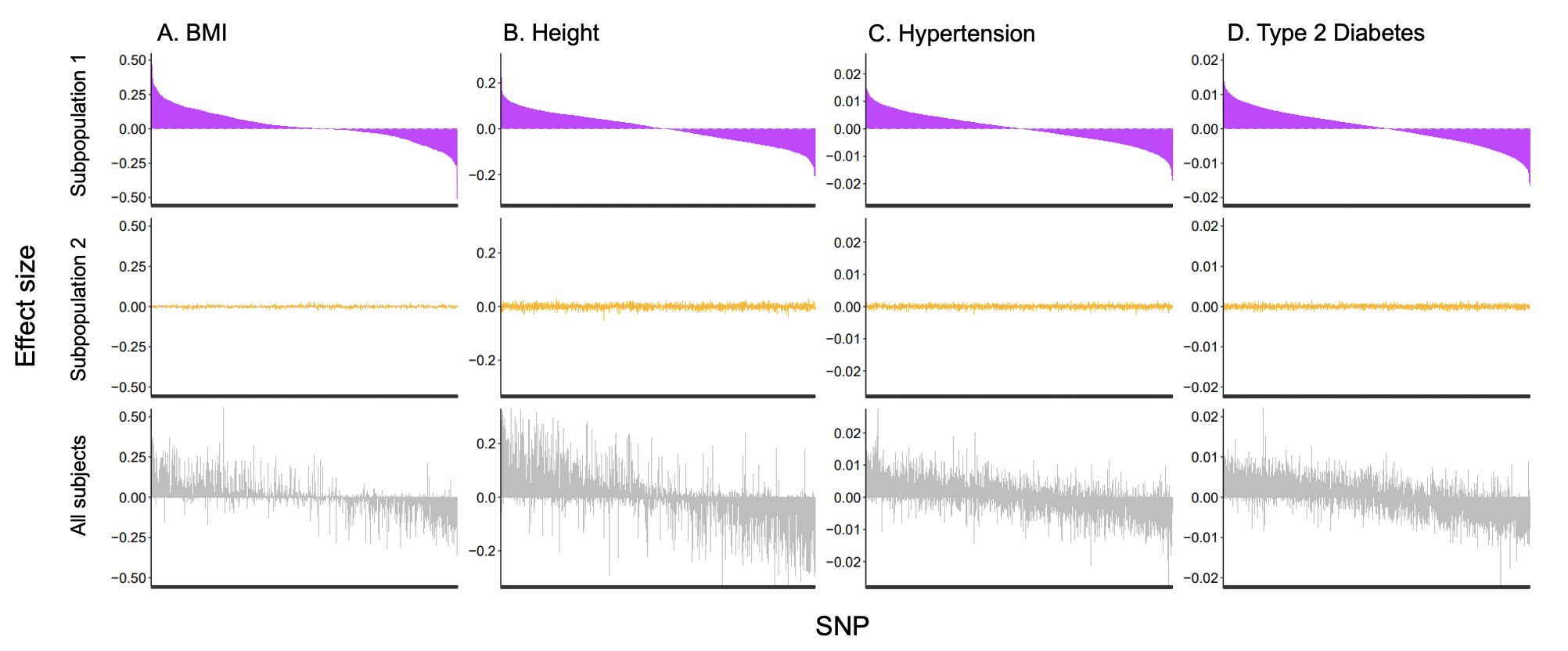

**Legend:** We selected top 5,000 significant SNPs in subpopulation 1 and estimate their effect sizes using DPR in the other subpopulations. In the horizontal axis, SNPs are ranked by effect size estimated in subpopulation 1.

**S12. Effect size stratification by subpopulations in UK Biobank data.**

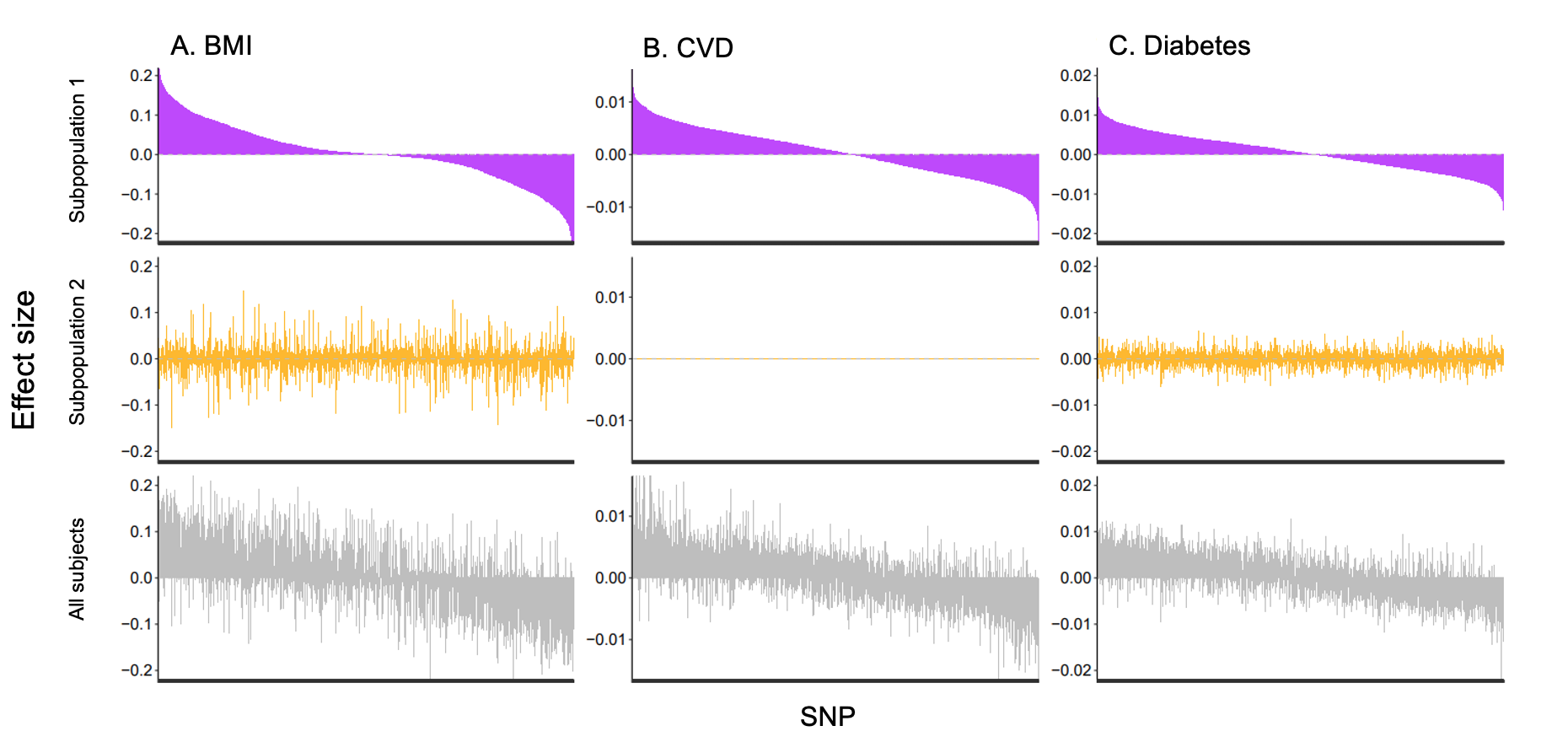

**Legend:** We selected top 5,000 significant SNPs in subpopulation 1 and estimate their effect sizes using DPR in the other subpopulations. In the horizontal axis, SNPs are ranked by effect size estimated in subpopulation 1. For CVD (panel B), the estimated effect sizes in subpopulation 2 is around the magnitude of ${10}^{-5}$ .

**Supplementary Notes**

**Simulation Study IV: The concordance of PV probability with ADMIXTURE**

In this simulation study, we aim to test whether it is concordant with exiting work to inferring individual genetic ancestry using PV framework. Referencing real GWAS data of European and African populations (dbGaP accession number: phs000021.v2.p1), an admixture population and two non-admixed “pure” ancestry cohorts consisting of 6,000 subjects and 10,000 SNPs were simulated (**Supplementary S13**). We calculated the PV probability of admixture population subjects assuming the two non-admixed cohorts are the two training strata. The calculated PV probability ${\Pr\left( \boldsymbol{x}_{s}|s\in k \right)}/{\sum_{k} \Pr\left( \boldsymbol{x}_{s}|s\in k \right)}$ matched closely to the genetic ancestry fraction estimated by ADMIXTURE (23) with Pearson correlation = 96.44% (**Supplementary S14**), which estimates the ancestry fraction based on direct genotype data.

**S13: The genetic ancestries in simulation study IV.**

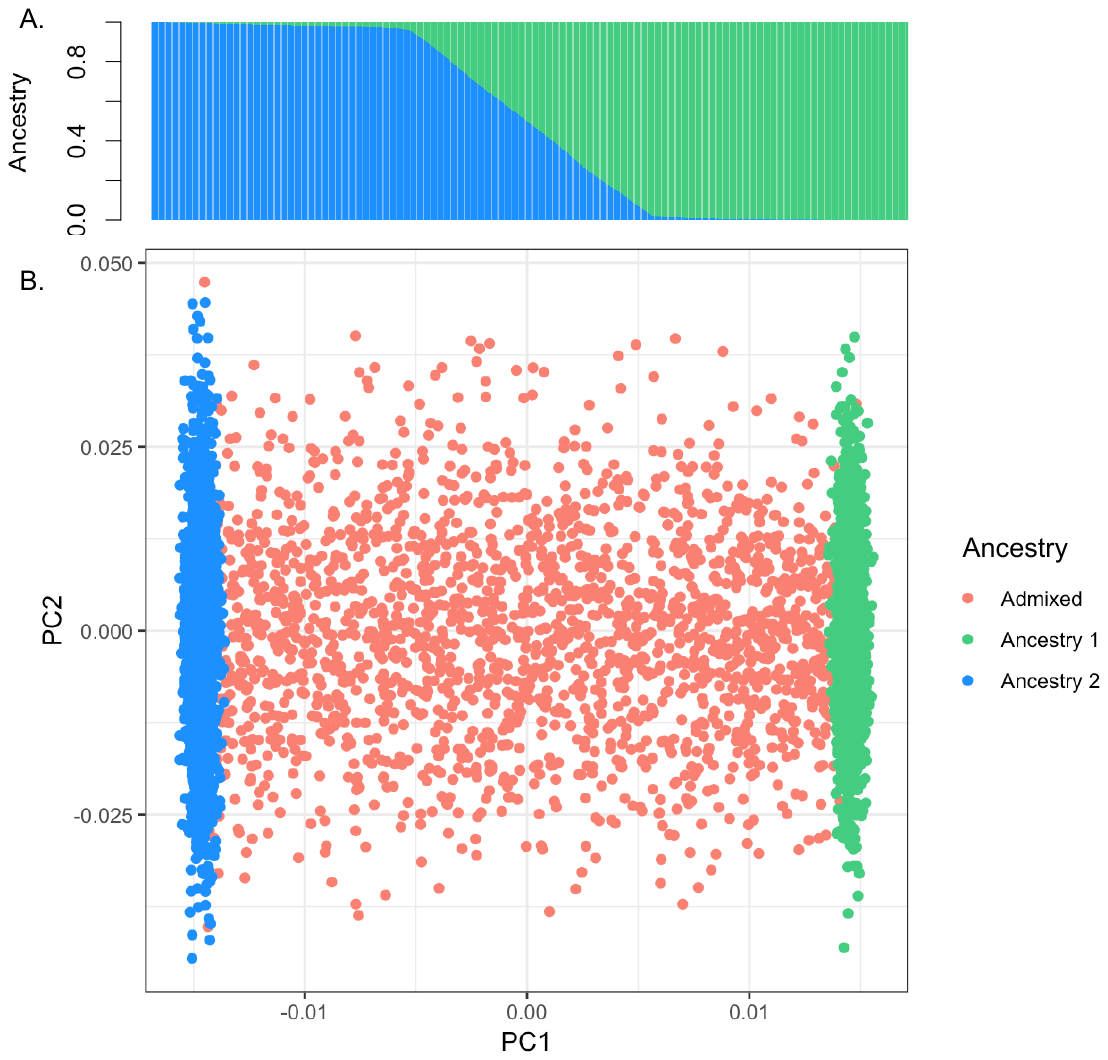

**Legend:** A. The inferred ancestries of the simulated admixture population. B. The admixed population plotted in the coordinates spanned by the top two principal components (PCs).

**S14. The comparison of PV and ADMIXTURE in calculating the genetic fraction.**

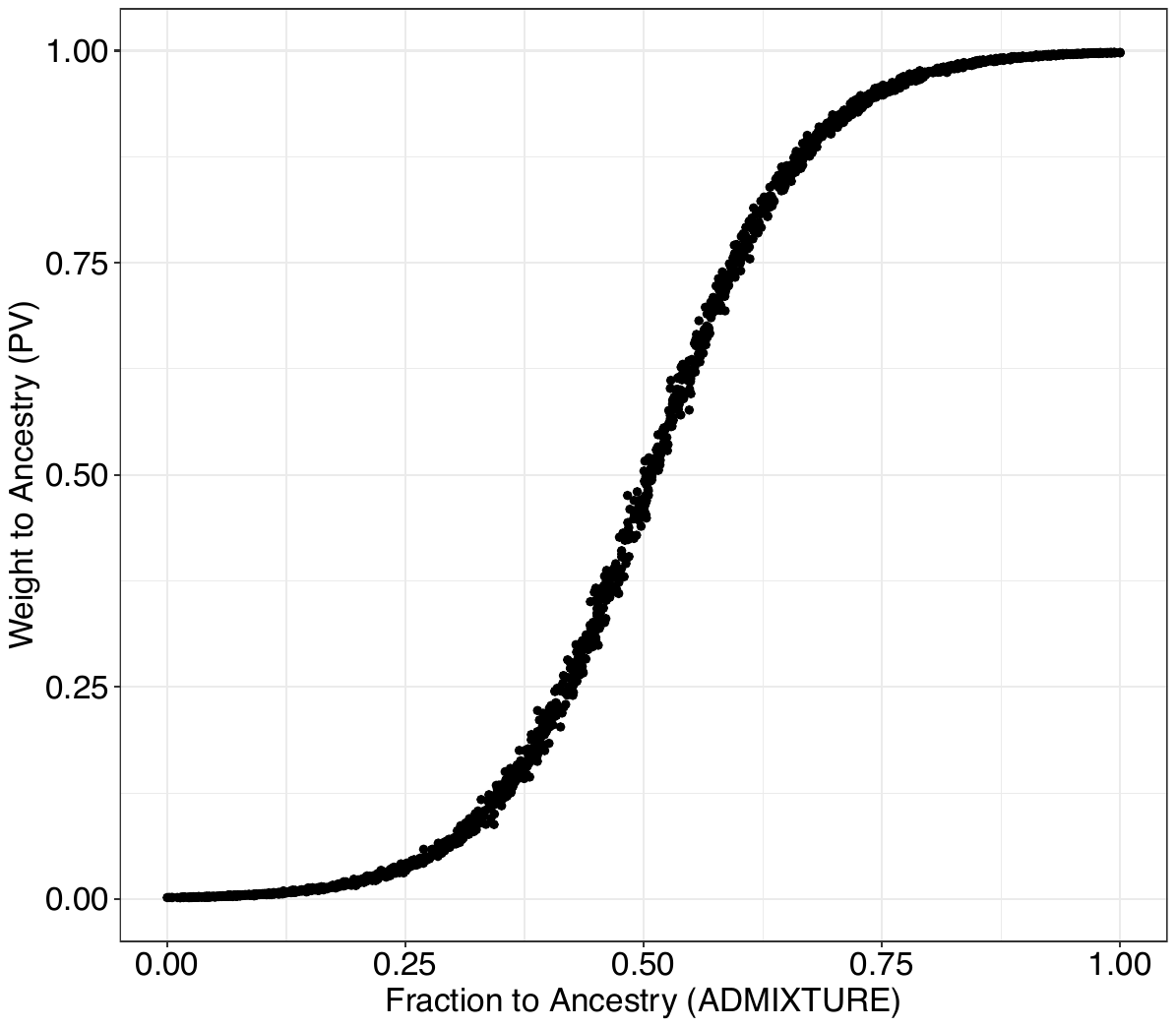

**Legend**: PV and ADMIXTURE are both applied to simulation data 1 to calculate the genetic ancestry proportion of to the admixed samples pure population 1. Top 10 eigenvectors were used in PV. In ADMIXTURE, K was set to 2.
